# Human Als3p Antibodies are Surrogate Markers of NDV-3A Vaccine Efficacy Against Recurrent Vulvovaginal Candidiasis

**DOI:** 10.1101/324483

**Authors:** Priya Uppuluri, Shakti Singh, Abdullah Alqarihi, Clint S. Schmidt, John P. Hennessey, Michael R. Yeaman, Scott G. Filler, John E. Edwards, Ashraf S. Ibrahim

## Abstract

A Phase 1b/2a clinical trial of NDV-3A vaccine containing a Candida albicans recombinant Als3 protein protected women <40 years old from recurrent vulvovaginal candidiasis (RVVC). We investigated the potential use of anti-Als3p sera as surrogate marker of NDV-3A efficacy. Pre- and post-vaccination sera from subjects who experienced recurrence of VVC (R) versus those who were recurrence-free (non-recurrent, NR) were evaluated. Anti-Als3p antisera obtained were evaluated for; 1) titer and subclass profile; 2) their ability to influence C. albicans virulence traits including hyphal elongation, adherence to plastic, invasion of vaginal epithelial cells, biofilm formation on plastic and catheter material, and susceptibility to neutrophil killing in vitro. Serum IgG titers in NR patients were consistently higher than in R patients, particularly for anti-Als3 subclass IgG2. Sera from vaccinated NR patients reduced hyphal elongation, adhesion to plastic, invasion of vaginal epithelial cells and biofilm formation significantly more than pre-immune sera, or sera from R- or placebo-group subjects. Pre-adsorption of sera with C. albicans germ tubes eliminated these effects, while heat inactivation did not. Finally, sera from NR subjects enhanced neutrophil-mediated killing of C. albicans relative to pre-immune sera or sera from R patients. Our results suggest that higher Als3p antibody titers are associated with protection from RVVC, attenuate C. albicans virulence and augment immune clearance of the fungus in vitro. Thus, Als3p serum IgG antibodies are likely useful markers of efficacy in RVVC patients vaccinated with NDV-3A.

**Abbreviations:**

Als3p: Agglutinin-like sequence 53 3 protein
AUC: area under the curve
CFU: colony forming unit
ConA: Concanavalin A
ELISA: enzyme-linked immunosorbent assay
Hyr1p: hyphal regulating protein 1
IRB: institutional review board
OPK: opsonophagocytic killing
NR: non-recurrent
NDV-3: recombinant His-tagged N-terminus of Als3p R formulated with alum
NDV-3A: recombinant N-terminus of Als3p R formulated with alum recurrent
RVVC: recurrent vulvovaginal candidiasis
ROC: Receiver-operating characteristic
Sap2: secreted aspartyl proteinase 2
SE: silicone elastomer
VVC: vulvovaginal candidiasis
YNB: yeast nitrogen base
YPD: yeast peptone dextrose

## INTRODUCTION

*Candida* species cause distressing mucocutaneous infections of the integument, oral and genitourinary tracts. Vulvovaginal candidiasis is estimated to occur in 50-75% of women in their childbearing years (1–3) and recurrence of vulvovaginal candidiasis (RVVC) is common (4). Hematogenously disseminated candidiasis is a life-threatening condition of increasing incidence in recent decades (1). Despite the use of antifungal therapy, candidemia is associated with ~40% attributable mortality (5). Compounding these concerns is the alarming rise in emergence of *Candida* species resistant to antifungal drugs (6).

*C. albicans* has multiple putative virulence capabilities including avid adherence to abiotic and host surfaces (7), the capacity to produce tissue-invading filaments (hyphae) (8), and the development of biofilms that promote immune evasion and impede efficacy of antifungal therapy (6). Targeting of these key virulence mechanisms provides opportunities for developing novel therapeutic interventions with minimal effects on the host mycobiome, and reduction in selection pressures that favor drug resistance (9).

NDV-3 is a vaccine containing a His-tagged recombinant version of the *C. albicans* Als3 protein (Als3p) N-terminus formulated with alum. Expressed on *C. albicans* hyphae, Als3p promotes adhesion of the fungus to biotic and abiotic substrates, enables invasion of host cell tissues, and facilitates biofilm formation (10, 11). Deletion of the *Als3* gene significantly impairs these virulence traits of *C. albicans in vitro* (10, 11). Consistent with these themes, NDV-3 decreases disease severity caused by *Candida* species in mice (12–15).

A Phase 1 clinical trial in healthy adults demonstrated safety and immunogenicity of the NDV-3 vaccine as evidenced by robust antibody and T-cell immune responses (13). Furthermore, a single dose of NDV-3A (rAls3p without the His-tag and formulated with alum) administered intramuscularly was safe and induced strong antibody and T-cell immune responses in patients with RVVC in a recent exploratory Phase 1b/2a study. This immune response protected patients <40 years of age with a history of RVVC from recurrence over a twelve-month study period (16). Specifically, post-hoc exploratory analysis revealed a statistically significant increase in the percent of the symptoms-free patients at twelve months post vaccination (42% vaccinated vs. 22% placebo; p=0.029) and a doubling time to first symptomatic episode (210 days vaccinated vs. 105 days placebo) for the subset for the patients <40 years of age (n=137).

The objective of the current study is to investigate the role of Als3p antibodies induced by NDV-3A as biomarkers of vaccine efficacy by quantitative and qualitative analysis of antibody titers and by evaluating the effect of these antibodies on *C. albicans* virulence traits.

## MATERIALS AND METHODS

### Serum Samples

All sera used in this study were prepared from blood collected from NDV-3A or placebo recipients in a Phase 1b/2a study in women with RVVC (ClinicalTrials.gov access number, NCT01926028) (16) using previously described methods (13) and were stored at −80°C until analyzed. Sera were obtained from 64 of 66 NDV3-A recipients and 53 of 60 placebo recipients using appropriate collection, processing and storage practices. In the NDV3-A group, 27 patients had no recurrence of VVC during the 12-month follow-up period and were classified as “non-recurrent” (NR), while 37 patients had one or more recurrences of VVC and were designated “recurrent” (R). For the placebo group, only 7 patients were classified as NR, while the rest were classified as R. Because of the low number of NR patients in the placebo arm all comparisons among NR and R patients were confined to NDV-3A vaccinated subjects. For the *in vitro* studies, matched sera from pre-immune (day 0) and post-vaccination (day 14 or 28) patients or placebo control were compared.

### *Candida* Strain

*C. albicans* SC5314 is a well characterized strain, and was the source the N-terminus of Als3 used to develop the NDV-3A vaccine (17). Routinely, the organism was cultured overnight in yeast peptone dextrose (YPD) broth (Difco) at 30°C with shaking prior to use for *in vitro* assays. To induce germination, *C. albicans* blastospores (5 × 10^6^) were grown in RPMI 1640 with L-glutamine (Gibco BRL) for 1 h at 37°C. For *in vivo* studies, *C. albicans* was serially passaged overnight 3 times in YPD before challenge in mice. In all studies, *C. albicans* cells were washed twice with endotoxin-free Dulbecco’s PBS, suspended in PBS or yeast nitrogen base (YNB, Difco) and counted with a hemocytometer to prepare the final inoculum.

### Analysis of Sera Components

Als3p antibody titers in sera were measured using an ELISA assay as previously described (13). To inactivate complement, aliquots of patient sera were independently heated at 55°C for 1 h, added to wells containing *C. albicans* in Yeast Nitrogen Base (YNB) medium, and incubated for 24 h at 37°C to permit biofilm development. To adsorb anti-*C. albicans* antibodies, the sera were incubated with *C. albicans* germ tubes for 1 h with gentle shaking at room temperature. The mixture was centrifuged at 21,000 *g* prior to using the cell-free supernatant in the biofilm assay. The presence and/or extent of removal of anti-Als3 antibodies (total IgG, IgG1 and/or IgG2) was measured by ELISA (18).

### PBMC Analysis

Peripheral blood mononuclear cells (PBMC) were collected from vaccinees as previously described (13). PBMCs were evaluated by ELISpot analysis to determine the portion of cells that could be stimulated to produce interferon (IFN)-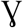 and IL-17A. Results are expressed in spots forming units (SFU) per 10^6^ cells.

### Adhesion and Biofilm Assays

Adhesion and biofilm formation were measured in 96-well polystyrene microtiter plates as previously described (19). Briefly, a 95 μL of *C. albicans* blastospores (2×10^5^ cells/ml in YNB medium) was added to wells containing 5 μL of patient serum (5% serum vol/vol), and incubated at 37°C. Control wells had no serum. After 2 h, wells were washed twice with PBS and the extent of adhesion was quantified by XTT assay (490 nm) (20). In parallel, cells were grown in the presence of 5% serum for 24 h to promote biofilm formation. Biofilms were washed twice prior to examining by bright field microscopy and quantification by XTT assay (20). Formation of biofilm on the catheter material silicone elastomer (SE) was also assessed (19). Briefly, circular SE pieces were pre-incubated with fetal bovine serum overnight at 25°C, washed twice and then introduced into the wells. The biofilm assay was conducted as above, in the presence or absence of patient sera.

### Invasion Assay

The human Ect1/E6E7 vaginal epithelial cell line was maintained in keratinocyte serum-free medium (Gibco) supplemented with bovine pituitary extract, epidermal growth factor, penicillin/streptomycin and passaged every 3-4 days as previously described (21). To study the effect of patient sera on *C. albicans* invasion, fibronectin-coated plastic coverslips were placed in a 24-well plate and the cells allowed to adhere overnight. After two washes, *C. albicans* cells were added to wells (fungus:host cell ratio of 5:1) for 12 h in the presence or absence of 5% patient serum. Non-adherent *C. albicans* was washed away, and the coverslips were stained with Concanavalin A (ConA) for 30 min at 37°C. The extent of epithelial cell invasion was visualized by differential staining using a confocal scanning laser microscope (Leica SP2) by overlaying the bright field image with a 594 nm excitation filter (red laser) for ConA. The non-invading yeast were stained, while the invading cells were unstained. The ability of *C. albicans* to invade the epithelium was expressed as % Invasion defined as: number of *C. albicans* cells invaded into the epithelium (i.e. unstained hyphae)/total number of *C. albicans* cells in a single bright field (stained + unstained cells) *100. At least 20 field per slide were blindly scored and presented as mean % invasion.

### Neutrophil Killing Assay

After obtaining IRB approved consent (LA Biomed protocol # 11672-07), neutrophils were isolated from blood collected from non-vaccinated human volunteers using endotoxin-free Ficoll-Paque Plus reagent (Amersham Biosciences) (12). Neutrophils were incubated with *C. albicans* germ-tubes containing YNB with 5% serum at 37°C without shaking (neutrophil:fungus ratio, 5:1). Controls contained *C. albicans* without neutrophils. After 90 min, the mixtures were sonicated to disrupt neutrophils and the surviving fungi quantitatively cultured. The percentage of opsonophagocytic killing (OPK) was calculated by dividing the number of CFU in the tubes containing neutrophils by the number of CFU in tubes without neutrophils.

### Statistical Analysis

All *in vitro* studies were performed in triplicate at a minimum, with two biological replicates. Different groups were compared using the non-parametric Wilcoxon rank sum test for pairwise comparisons, and Mann Whitney test for comparison of unmatched groups. Data were analyzed in GraphPad Prism software (LaJolla, CA), and a p-value < 0.05 was considered statistically significant. We estimated how well the *in vitro* assays of adhesion, biofilm formation, neutrophil killing and IgG2 titers discriminated between sera of R and NR patients. We performed Receiver-operating characteristic (ROC) analysis on GraphPad Prism, which visualizes the sensitivity and specificity characteristics of a particular assay. The y-axis of the ROC graph represents sensitivity, or the true positive rate, i.e. the proportion correctly discriminated or predicted as by the assays. The x-axis represents the component complement of specificity (100% - specificity). Area under the ROC curve (AUC) is a commonly used measure, where AUC of 1.0 represents a perfect curve fit, while an AUC of 0.5 represents random classification (22). Using ROC, we determined the cut point that maximized the sum of sensitivity and specificity for all the assays.

## RESULTS

### Antisera from R and NR Subjects Had Distinct Quantitative and Qualitative Antibody Profiles

We analyzed the antibody titers of sera from R or NR patient in an attempt to understand the partial protection elicited by NDV-3A vaccine. We conducted an area under the curve (AUC) analysis of the total IgG titers of sera collected from NDV-3A vaccinated subjects over the 12-month period. AUC of total IgG titers from NR patients was significantly higher than those in sera from R patients during the early time points of collection (Day 0-90) (Figure 1A). Similarly, the geometric mean of IgG titers of the NR patients at 14 or 28 days post vaccination was approximately twice as high as that detected in R patients (Figure S1 in Supplemental Material). At later time points (Days 90-360), and despite the general drop in Ab titers for both patient populations, the difference in AUC titers of the NR vs. R patients was even greater (Figure 1B). Importantly, while the vast majority of placebo patients had first recurrence within the first 90 days post vaccination (median recurrence of 53 days), most of the vaccinated subjects had their first recurrence later than this (median recurrence of 94 days) (Figure 1C). Consistent with the Phase 1b/2a clinical trial (16), the enhanced time to recurrence was significant among vaccinees who are <40 years old (p= 0.043). Interestingly, the later recurrence corresponded with the decreased IgG levels beyond 90 days.

**Figure 1.**
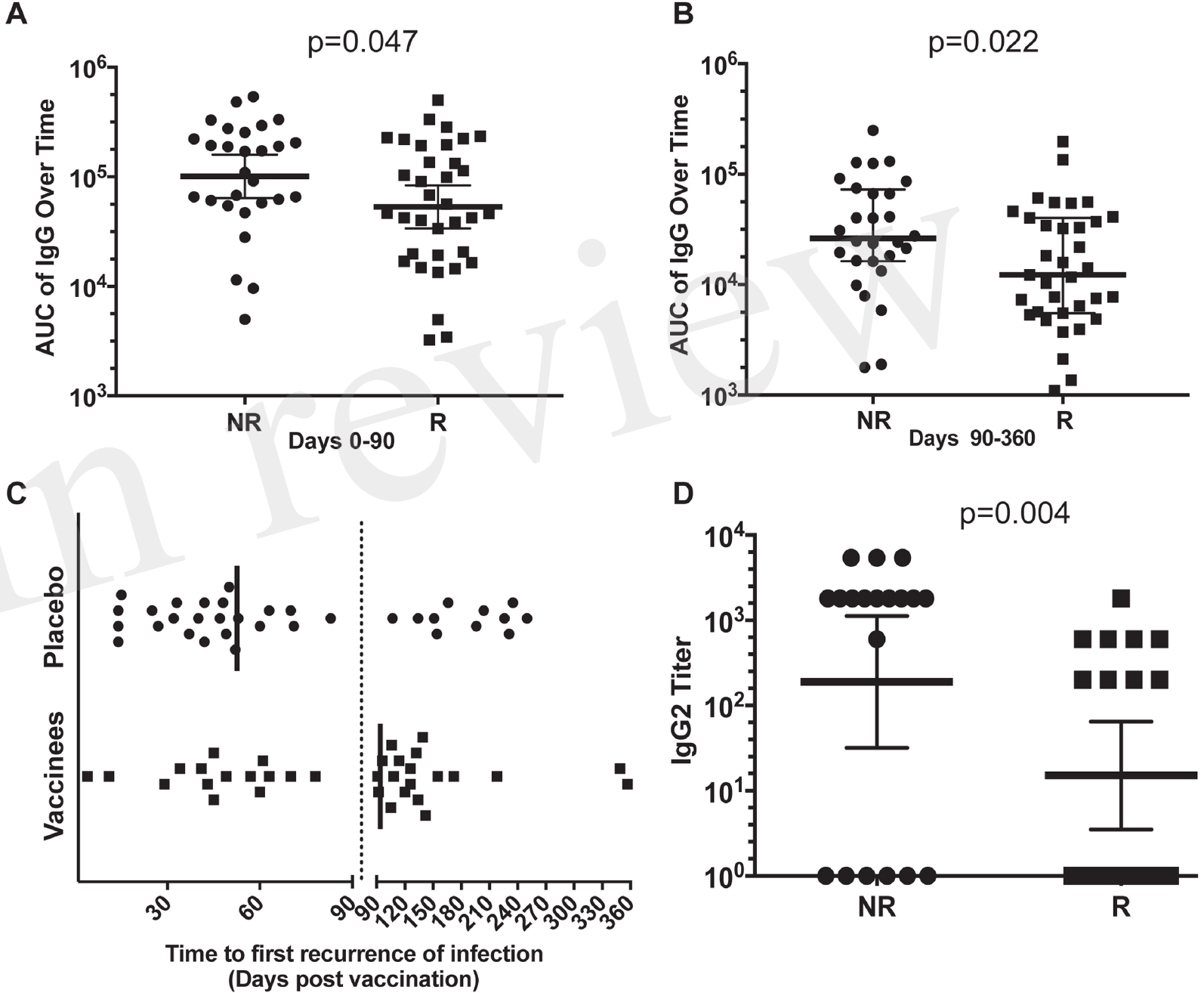
Analysis of the antibody response in vaccinated patients. Mean area under the curve (AUC) of the total IgG titers over time (0-90 and 90-360 days) for each patient, in the NR and R NDV-3A-vaccinated subjects was plotted. In the first 3 months post-vaccination, AUC of NR patients was significantly higher (p=0.046) than that of R patients (A). In months 3 – 12, this difference in AUC was also significant (p=0.022) (B). The decrease in AUC of the IgG titers in R patient sera in the later months corresponded with the increase in recurrent episodes of VVC during this period (C). Finally, significantly more number of NR patients displayed IgG2 antibodies in their sera, also mean (geometric) IgG2 antibody titer was higher in NR patients, compared to R patients (p=0.003) (D). Each dot in A, B, and D represents antibody titers in each analyzed serum samples from the indicated individual patients. Each dot in C represent a first relapse in infection as a function of time post vaccination. Data in A, B, and D are presented as geometric mean with 95% confidence interval.

In parallel, the serum antibody profiles were evaluated for anti-Als3 IgG subclasses. The IgG1 subclass comprised the predominant isotype in vaccinated NR and R sera (NR vs R IgG1 titer, p=0.9, data not shown). Remarkably, the IgG2 titer in NR sera was much higher than in the R patient sera (Figure 1D), suggesting an isotype-specific enrichment in NR immune responses. Titers of IgG 3 and IgG4 were not significantly different in sera of R vs. NR patients (data not shown). Also, we did not find any differences among IFN-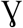 or IL-17 levels between R vs. NR patients, despite the enhanced level of these two cytokines upon vaccination with NDV-3A (Figure S2 in Supplemental Material). The quantitative and qualitative differences among serum antibodies from R vs. NR patients prompted us to test their effect on *C. albicans* virulence traits below.

### Sera from NR Subjects Reduced *C. albicans* Adhesion to Plastic

Post-vaccination sera from NR subjects that received NDV-3A significantly reduced adhesion of *C. albicans* to plastic, compared to pre-vaccination sera from the same patients (Figure 2A). In contrast, post- vaccination sera obtained from R patients that received NDV-3A did not significantly alter *C. albicans* adhesion relative to pre-vaccination sera (Figure 2B). As expected, sera from patients who received the placebo did not influence *C. albicans* adhesion to plastic (Figure 2C). Sera from the NR-NDV-3A cohort was the only one that significantly reduced adhesion, compared to the other two groups (Figure 2D).

**Figure 2.**
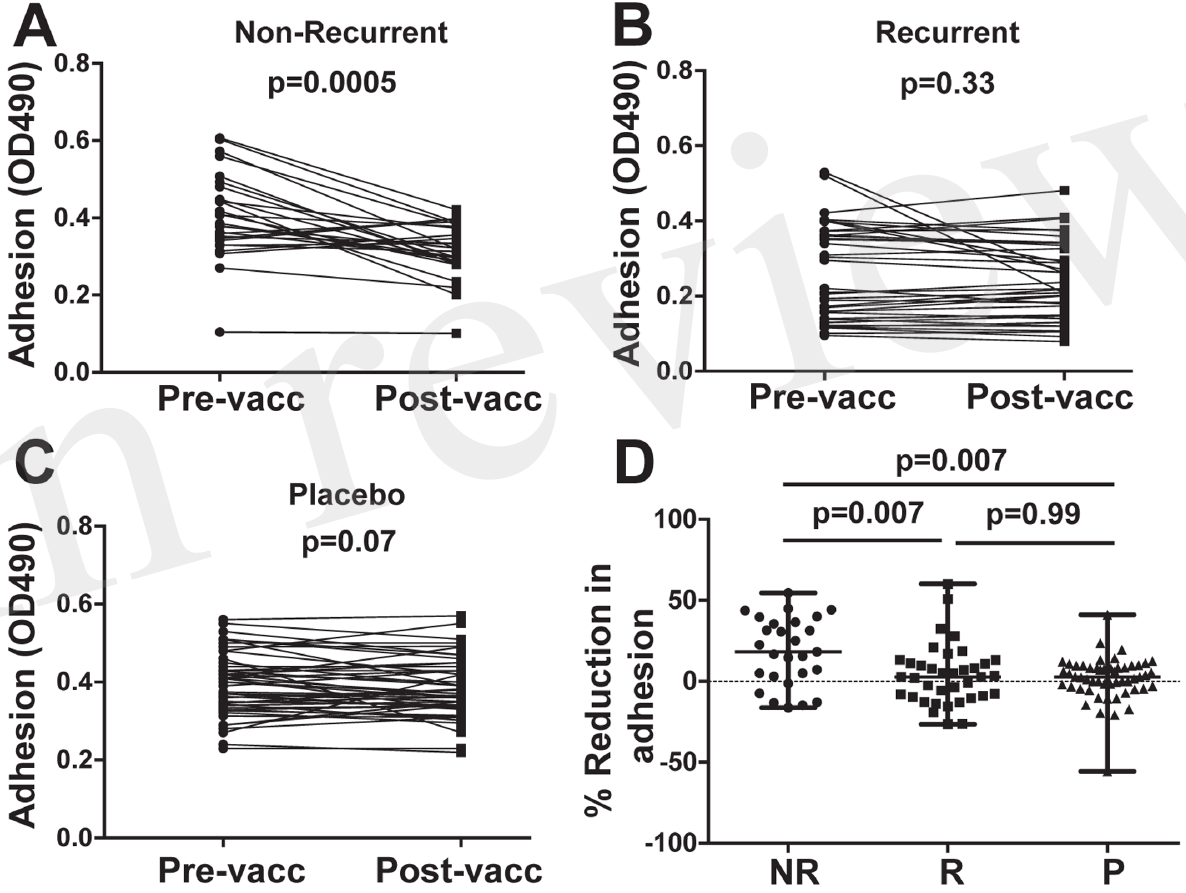
*In vitro* assessment of *Candida albicans* adherence to plastic in presence of patient sera. Postvaccination sera from 27 non-recurrent NR patients significantly (p=0.0005) reduced *C. albicans* adhesion when compared to their respective pre-vaccination sera (A). There was no difference in the extent of adhesion between pre and post vaccination sera from 37 recurrent (R) patients (p=0.33) (B), or 53 placebo (P) patients (p=0.067) (C). Percent inhibition of *C. albicans* adhesion to plastic was significantly higher in post-vaccination sera of NR patients versus that of R or P patients (D). Data in D are presented as median ± interquartile range. Each dot represents alteration in *C. albicans* adhesion due to an individual patient sample.

### Sera from NR Patients Reduced *C. albicans* Biofilm Development

Post-vaccination sera from NR subjects who received NDV-3A reduced biofilm development as compared to their paired pre-vaccination sera (Figure 3A). This reduction was not observed when cells were incubated with sera from R subjects who received NDV-3A or sera from placebo recipients (Figure 3B, C). As in the adhesion assay, a significantly greater reduction in biofilm formation was observed in sera from NR versus R patients that received NDV-3A or placebo recipients (Figure 3D).

**Figure 3.**
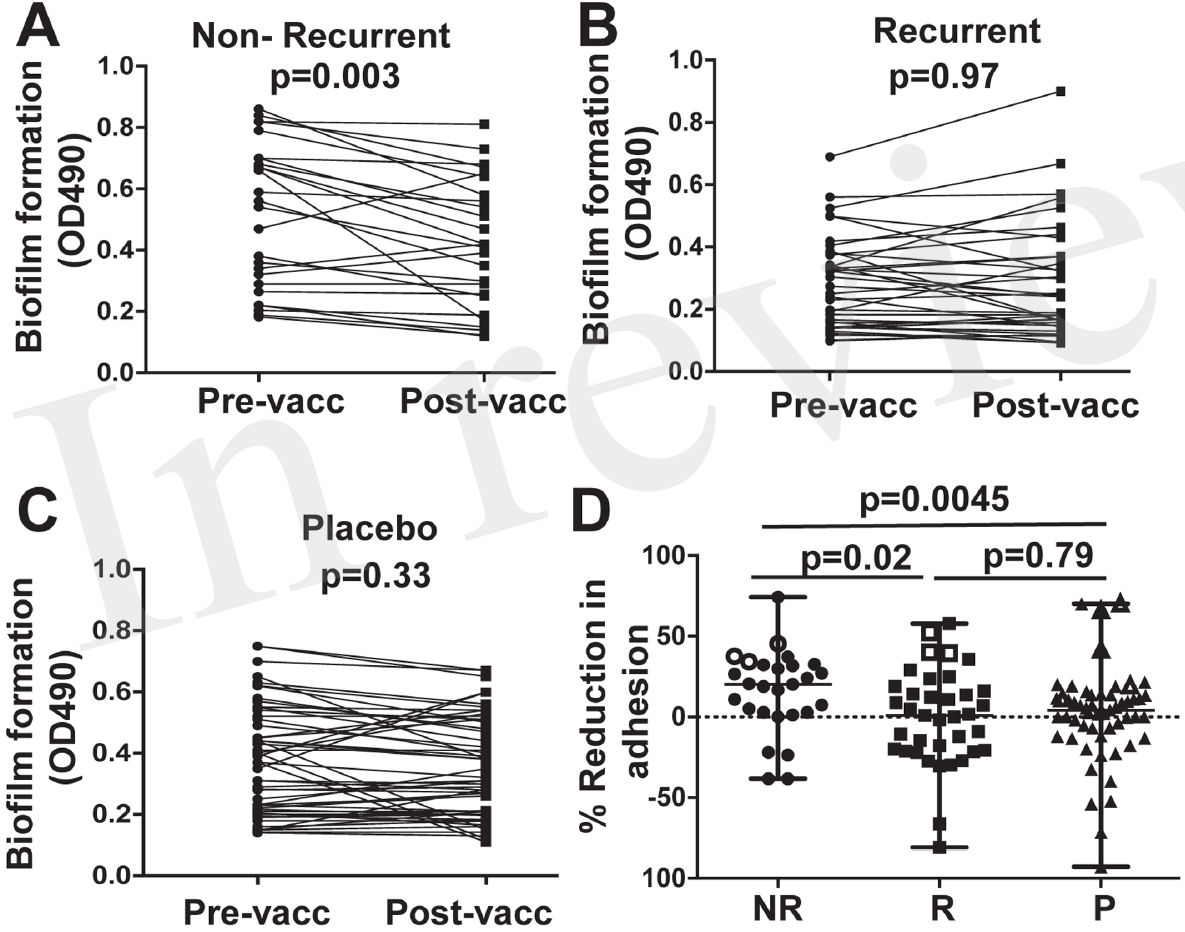
*In vitro* assessment of *Candida albicans* biofilm formation in presence of patient sera. Postvaccination sera from 27 non-recurrent (NR) patients significantly (p=0.003) reduced *C. albicans* biofilm formation on 96-well microtiter plates, when compared to their respective pre-vaccination sera (A). There was no difference in the extent of biofilm growth between pre- and post-vaccination sera from 37 recurrent (R) patients (p=0.97) (B), or 53 placebo (P) patients (p=0.33) (C). The percent inhibition of *C. albicans* biofilms to plastic was significantly higher in post-vaccination sera of NR patients versus that from R or P patients (D). Data in D are presented as median ± interquartile range. Each dot represents alteration in *C. albicans* biofilm formation due to an individual patient sample. The open data points in D represent 6 placebo and 6 NDV3-A vaccinated patients (3 R and 3 NR) whose sera showed the highest reduction in biofilm formation, and were chosen for the vaginal epithelial cell invasion assay presented in Figure 3B.

Bare-plastic is the gold-standard to measure biofilm formation (20). We wanted to confirm that sera samples that prevented biofilm formation on bare-plastic also prevent biofilm formation on SE used in manufacturing catheters. Thus, antisera of NR patients displaying the highest extent of biofilm inhibition were tested for their ability to impede biofilm formation on SE. These sera reduced *C. albicans* biofilm formation on the SE substrate to an extent similar to that observed in 96 well-plates (Figure S3 in Supplemental Material).

Consistent with reduction in biofilm formation, wells containing post-vaccination sera from NR patients also displayed reduced *C. albicans* adhesion by bright field microscopy, as depicted by reduced density of cells in the bottom of the wells (Figure 4A). This reduction in adhesion to plastic was accompanied by reduced hyphal elongation, but not germ tube induction (Figure S4 in Supplemental Material). In contrast, *C. albicans* growing in wells containing pre-vaccination sera from the same patient or commercially obtained pooled human serum (controls) displayed robust filamentation and biofilm formation (Figure 4A).

**Figure 4.**
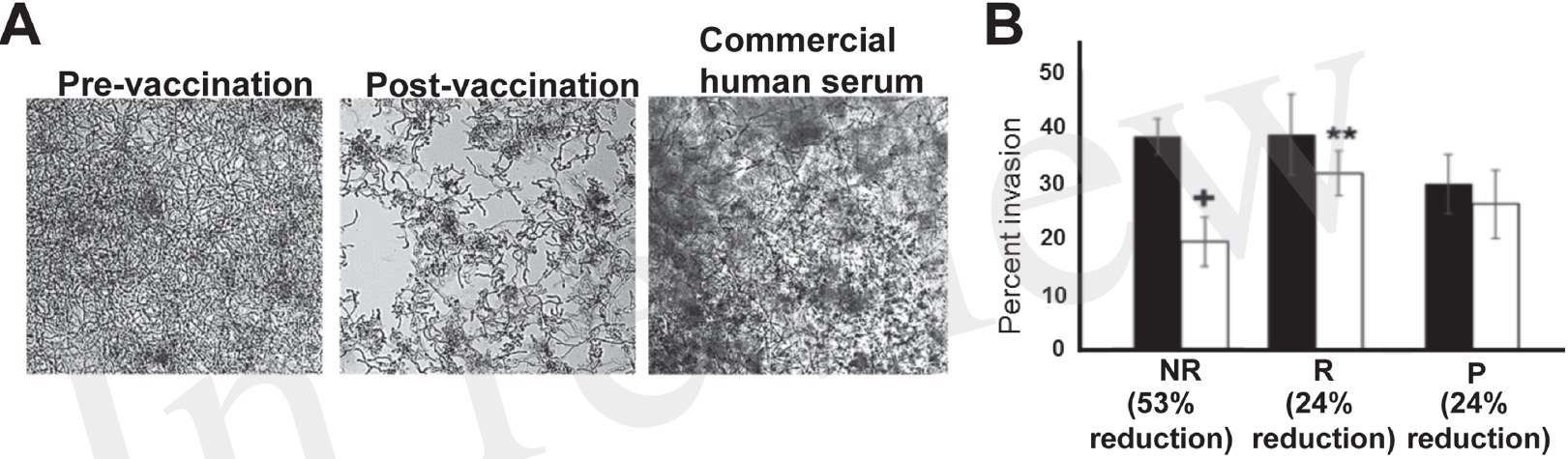
*In vitro* assessment of *Candida albicans* filamentation, and invasion of vaginal epithelial cells in presence of patient sera. Representative micrographs (from 6 different samples in each arm) showing that post-vaccination sera from NR patients that abrogated biofilm formation displayed short and wavy hyphae, compared to the normal robust hyphae in the biofilms formed in pre-vaccination serum, or the control commercial pooled human serum (A). Six samples from placebo or NDV-3A vaccinated patients (3 R and 3 NR) were selected from Figure 2D (open symbols) for analysis in invasion of vaginal epithelial cells. Post vaccination sera from NR patients inhibited invasion of vaginal epithelial cells two-fold more than R patient or P patient sera (B). **P <0.01 for post- vs. pre-vaccination sera from R patients. ^+^P <0.05 for post-vaccination sera vs. pre-vaccination sera from NR and vs. post-vaccination sera from R or placebo patients. Data in B are presented as mean ± SD.

### Sera from NR Patients Reduced *C. albicans* Invasion of Vaginal Epithelial Cells

Although sera preventing adhesion and biofilm formation were predomainantly from the NR group, some R and placebo subject sera also impeded these *C. albicans* virulence functions. Therefore, sera from the three patients exhibiting the greatest inhibitory effect in each group were compared for pre- and post-vaccination impact on *C. albicans* capacity to invade vaginal epithelial cells (Figure S5 in Supplemental Material). Antisera from NR patients reduced *C. albicans* invasion of epithelial cells by ~53%, approximately two-fold higher than inhibition displayed by antisera of R patients (24%) (Figure 4B). The sera from placebo patients did not inhibit *Candida* invasion in this assay (Figure 4B).

### The Inhibitory Function of Serum was Independent of Complement and Likely Associated with Antibodies

To evaluate the possible role of complement, sera from 12 NR patients that received NDV-3A and had the highest decrease in biofilm development were heat-inactivated and then retested in the *C. albicans* biofilm assay. Inactivated sera retained biofilm inhibition equivalent to that of native sera (Figure 5A), indicating that complement does not play a significant role in the capacity of the sera to inhibit biofilm formation.

**Figure 5.**
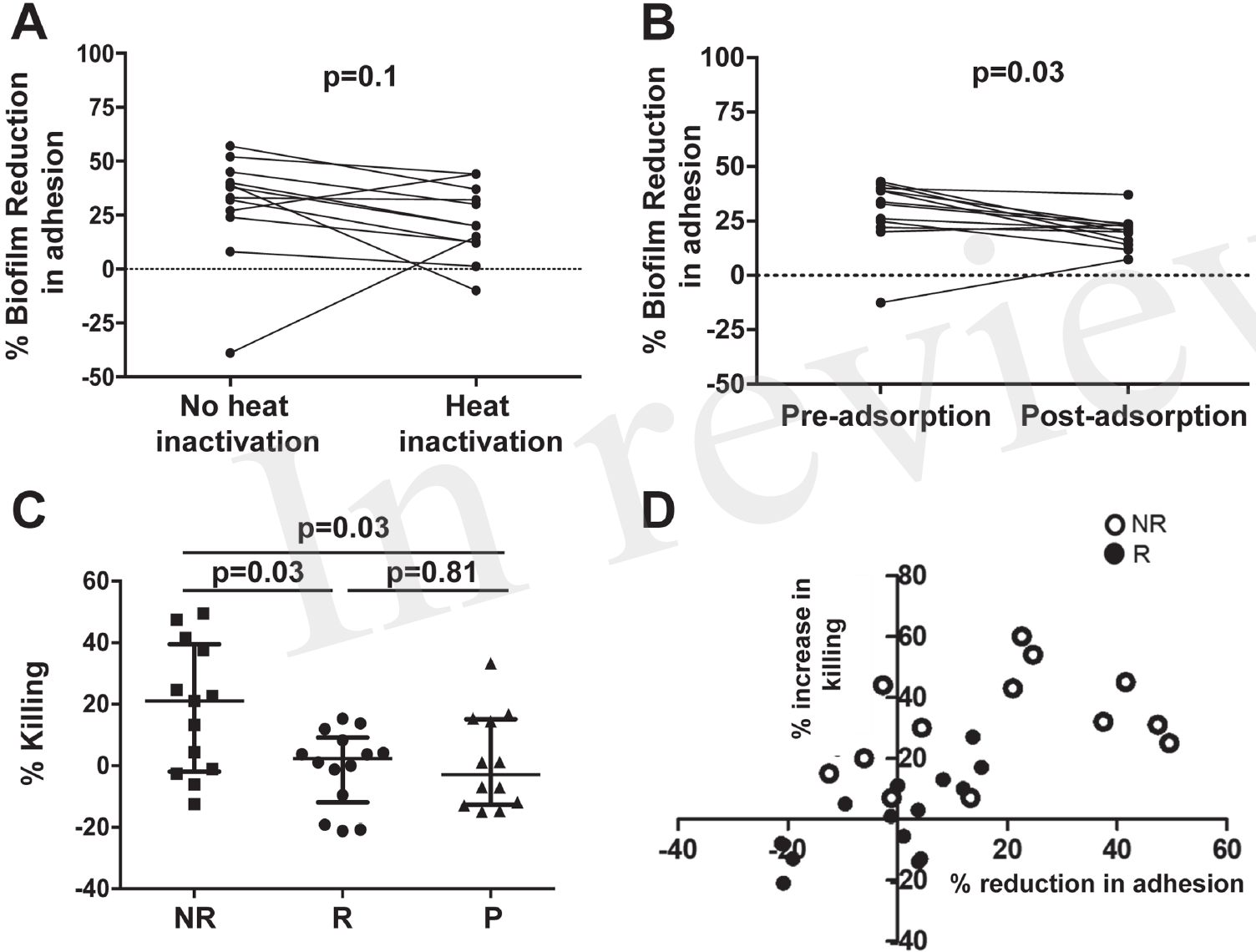
Assessment of the role of antibodies in affecting virulence, and evaluation of OPK of *C. albicans* germ tubes in the presence of patient sera. Heat treatment of post-vaccination sera from NR patients does not significantly reduce its biofilm-inhibitory activity, compared to paired untreated sera (A). Adsorption of antibodies in post-vaccination sera from NR patients with *C. albicans* germ tubes significantly (p=0.03) abolishes the biofilm inhibitory activity of the sera, when compared to paired adsorbed pre-vaccination sera. Only post-vaccination sera from NR patients significantly (p=0.03) enhance OPK and killing of *C. albicans* germ tubes by human neutrophils, compared to post-vaccination sera from R or P patients (C). A comparison between percent increase in OPK activity and percent reduction in adhesion, in post-vaccination sera from NR and R patients, resulted in significant correlation within the respective subject sera (D). Each open circle represents individual NR sera, which displayed both an overall greater reduction in adhesion and increase in neutrophil killing. Solid circles denote the individual R patients that compared to the NR patients, show a smaller % decrease in adhesion as well as neutrophil killing. Negative values on the graph represent % increase in adhesion or % decrease in neutrophil killing.

To determine whether anti-*C. albicans* antibodies were the active constituent of serum, we incubated the pre- and post-sera with *C. albicans* germ tubes to adsorb antibodies against Als3. This process will adsorb antibodies targeting all surface proteins that are expressed on *C. albicans* germ tubes including those targeting Als3p. Indeed, ELISA plates coated with rAls3p for both day 0 and post-vaccination confirmed that the absorption process significantly reduced the anti-Als3 IgG titers in the samples (Figure S6 in Supplemental Material). Next, these sera were used in *C. albicans* biofilm assays as detailed above. Adsorption of antibodies from post-vaccination serum reduced their ability to inhibit biofilm formation (Figure 5B).

### Sera from NR Patients Enhanced Neutrophil-Mediated Killing of *C. albicans*

We questioned whether such functionally active sera influenced interactions of the fungus with neutrophils from unvaccinated human volunteers *ex vivo.* As displayed in Figure 5C, sera from NR patients enhanced neutrophil killing of fungal cells, compared to sera from R or P patients.

We also determined whether sera from NR patients that demonstrated the highest reduction in *C. albicans* adhesion to plastic also exhibited the highest level of neutrophil-mediated killing. We found a strong correlation between the ability of sera from NR patients to reduce adhesion and increased neutrophil-mediated killing (p<0.05 and R^2^ of 0.66). Further, overall larger numbers of NR patients (13 NR subjects) induced OPK and prevented adhesion than the R group (13 subjects) (Figure 5D and Figure S7 in Supplemental Material).

### ROC Analysis of the *in vitro* Assays Predicts Biomarkers of Vaccine Efficacy

To statistically further validate the sensitivity and specificity of each of the *in vitro* assays, we performed an area under the ROC curve (cvAUROC) analysis for four assays comparing R to NR patient sera for patients that received NDV-3A. Our analysis of IgG2 predict that NR patients have higher IgG2 antibodies than 75% of the R patients (area under curve 0.75, p value 0.008). Further, our data reveal that an IgG2 antibody titer cutoff of above 1680 (100% sensitivity and >63% specificity) would predict the vaccinated patient to be protected. Similarly, a high sensitivity/specificity was obtained for the remaining three assays: adhesion (area under curve 0.7, p value= 0.007), biofilm reduction (area under curve 0.68, p value = 0.012), neutrophil killing (area under curve 0.75, p value= 0.029). These analyses support such correlations as potential biomarkers of vaccine efficacy.

## DISCUSSION

In this study, we had the unique opportunity to compare humoral immune responses in patients who derived a measurable health benefit from the NDV-3A vaccine versus those who did not, and versus placebo patients. In addition, sera from R vs. NR subjects could be compared for their ability to impede key virulence functions of *C. albicans in vitro*. The study goals were to explore potential surrogate biomarkers of protection that might be useful in future studies of this and more serious *Candida* infections and to gain insight into the potential mechanism(s) contributing to protective efficacy of the vaccine. Our studies focused on determining the impact of serum antibodies on selected putative virulence factors of *C. albicans* for three reasons: 1. The NDV-3 vaccine (based on *C. albicans* Als3 antigen with 6X His tag) induced high antibody titers (13); 2. High antibody titers predict NDV-3 vaccine efficacy in mice (23); and 3. Antibodies, including those targeting Hyr1p (24, 25), Sap2 (26), and *Candida* cell wall glycopeptides (27, 28), protect against experimental candidiasis. We focused also on *Candida* adhesion and invasion of vaginal epithelial cells, as well as biofilm formation, since Als3p is a known mediator of these putative virulence functions (10, 11, 29).

NR patients maintained a high antibody median titer (AUC) of ≥25,000, while R patients had a median AUC titer of 10,000 after 90 days post vaccination. This temporal waning of antibody response coincided with the increased frequency to first relapse in R patients. The decrease in antibody titer raises the possibility that R patients may benefit from a booster dose following priming with the vaccine. There was also a significant increase in IgG2 subclass titers in NR patients that received NDV-3A as compared to the R patients that received the vaccine, suggesting that this IgG subclass may be a surrogate marker of protection post-vaccination. Human IgG2 and IgG4, but not IgG1 or IgG3, have been reported to protect mice against *Cryptococcus neoformans* infection (30), most likely by enhancing the fungicidal activity of macrophages (31). Based on our current data, it is possible that the IgG2 subclass antibody component could impair *C. albicans* interactions with host tissues, and contribute to neutrophil activation leading to enhanced *C. albicans* killing by NR antisera. However, an important alternative hypothesis is also of interest, that IgG2 and IgG4 antibody are surrogates for non-inflammatory skewing of immune responses, biasing against symptoms of relapse.

Sera from NR patients that received NDV-3A significantly reduced *C. albicans* adherence to plastic and SE, and impeded invasion of vaginal epithelial cells by *C. albicans* hyphae more than antisera obtained from R patients that received NDV-3A or those from placebo. Biofilm formation is a function of the ability of *C. albicans* to adhere to abiotic surfaces. Thus, it was not unexpected that higher levels of anti-Als3 antibodies, as seen in antisera from NR patients but not R patients, would also significantly reduced *Candida* biofilm formation. As determined from antibody adsorption and complement inactivation studies, such abrogation of these *C. albicans* virulence functions was due, at least in part, to anti-Als3p antibodies and did not require complement fixation.

Our group previously demonstrated that NDV-3 protects mice from VVC by a mechanism that involves priming of both B cell- and T cell-mediated adaptive immune responses (12). Specifically, anti-Als3p antibodies enhanced the *ex vivo* killing of *C. albicans* by neutrophils primed with IFN-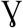 (12). Although in the current study we could not detect a correlation between IgG titers of NR and R vaccinated subjects and their corresponding IFN-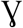 levels, antisera from NR patients significantly enhanced the ability of human neutrophils to kill *C. albicans ex vivo* as compared to antisera from R patients or patients administered placebo. These results are concordant with the finding that RVVC is a disease in which a discordance of exacerbated neutrophil influx often occurs in the face of inefficiency in clearing the infection (32–34). We postulate that in NR women, the vaccine was able to induce an antibody response that; 1) protected against *C. albicans* adherence to and invasion of mucocutaneous barriers; 2) reduced the capability of the organism to form biofilm from which persistent infection occurs; 3) induced a coordinated phagocyte response that is more efficacious in clearing the infection; and/or 4) modulated profusive inflammatory responses of the host associated with relapse.

The statistical robustness of the *in vitro* assays of IgG2, adhesion, biofilm formation and neutrophil killing which were validated in our ROC analyses, revealed that any of these tests could be used as biomarkers of vaccine efficacy. While the latter three assays are likely too cumbersome to support larger clinical trials, an ELISA measuring IgG2 in the sera of vaccinated patients would be a simple yet robust method to predict the protective efficacy of a vaccine in a human clinical study like the current one, or future studies on disseminated candidiasis.

Our forthcoming studies are planned to precisely define the roles of antibody isotypes, the impact of boosting, the combination of multiple antigen vaccines, and the influence of advanced adjuvants to further optimize vaccine and immunotherapeutic strategies targeting *Candida* species.

**Figure 6.**
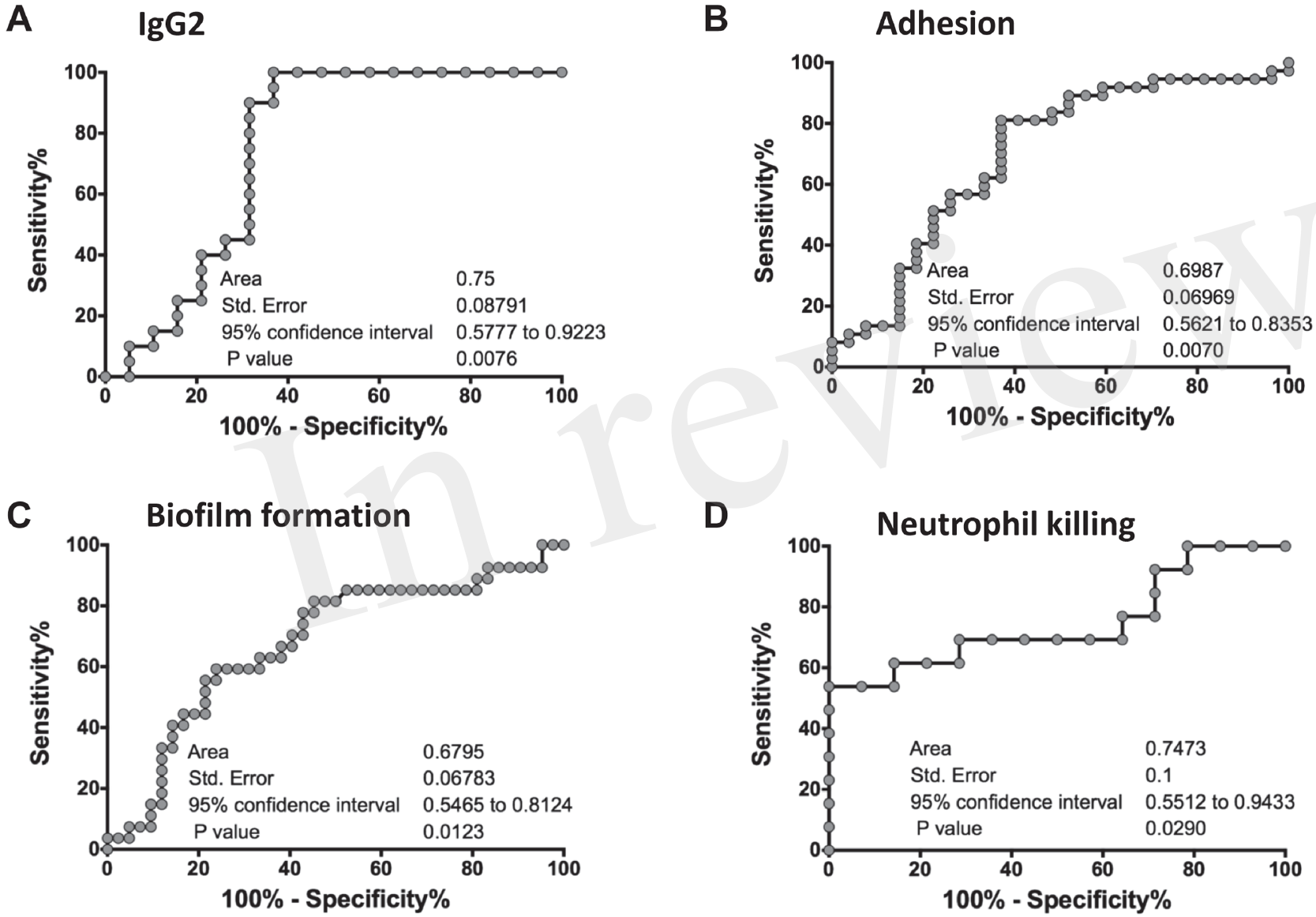
ROC analysis of the *in vitro* assays. An ROC analyses for four *in vitro* studies was performed on GraphPad Prism software, where a graph was generated of 100% - (minus) Specificity% versus Sensitivity % for each of the assays: IgG2 titers (A), adhesion (B), biofilm formation (C) and neutrophil killing (D). For each graph, an Area Under the Curve (Area), standard error of the AUC under the ROC curve, as well as the 95% confidence interval is reported. A p-value of <0.05 in each of the ROC curves concludes that the results are significant, and robust.

## AUTHORS CONTRIBUTIONS

PU designed, performed, supervised the project and wrote the manuscript. SS performed experiments, designed the ROC, analyzed the data and revised the manuscript. AA performed the experiments and analyzed the data, CS provided materials and revised the manuscript. JH, provided materials and revised the manuscript, MY revised the manuscript, SF contributed to the study design and revised the manuscript. JE contributed to the study design and revised the manuscript. AI designed and supervised the project and wrote the manuscript.

## ETHICS STATEMENTS

This study was carried out in accordance with the recommendations of National Institutes of Health guidelines for human subject policies and ethical guidance and regulations with written informed consent from all subjects. All subjects gave written informed consent in accordance with the Declaration of Helsinki. The protocol was approved by the ‘Los Angeles Biomedical Research Institute IRB’.

